# Extracellular vesicles propagate aging in COPD airway epithelial cells by transfer of microRNA-34a

**DOI:** 10.1101/2023.12.18.572220

**Authors:** Justine V. Devulder, Jonathan R. Baker, Peter S. Fenwick, Lina Odqvist, Louise E. Donnelly, Peter J. Barnes

## Abstract

**Rationale:** Chronic obstructive pulmonary disease (COPD) is associated with the acceleration of lung aging, demonstrated by the accumulation of senescent cells in lung tissue. MicroRNA (miR)-34a is induces senescence by suppressing the key anti-aging molecule, sirtuin-1 (SIRT1). Senescent cells spread senescence to neighboring and distant cells, which favors the progression of COPD and its comorbidities. The mechanisms for spreading senescence remain undetermined but may be mediated by the transfer of microRNAs in extracellular vesicles.

**Objectives:** To analyze the miRNA content of extracellular vesicles in COPD and explore their effect on cellular senescence of healthy cells

**Methods:** EVs were isolated from small airway epithelial cells (SAEC) from healthy donors or COPD patients. Recipient healthy SAEC were cultured with EVs and the expression of miR-34a and markers of cellular senescence, p21^CIP1^ and SIRT1, were measured.

**Main Results:** EVs from COPD cells induce senescence in healthy recipient cells via the selective transfer of miR-34a. We showed that COPD SAEC produce increased numbers of EVs enriched with miR-34a. EVs are taken up by healthy cells, resulting in reduced expression of the anti-aging molecule sirtuin-1 and increased expression of markers of senescence, such as p21^CIP1^ and positive staining for senescence-associated β-galactosidase

**Conclusions:** Our findings provide evidence of the mechanism by which EVs spread cellular senescence in human primary cells via miR-34a, rather than via soluble mediators. EVs enriched with miR-34a may spread senescence locally, accounting for disease progression, but also provide a mechanism for distant spread to account for comorbidities and multimorbidity of the elderly.

## Introduction

Chronic obstructive pulmonary disease (COPD) is a condition of accelerated lung aging (1). Indeed, aging is associated with increased vulnerabilities to chronic diseases and multimorbidity (2, 3) with major consequences on the burden of health care (2). Aging is a major risk factor associated with COPD and its prevalence is two to three times higher in patients over 60 years than in younger age groups (4). Cellular senescence contributes to the aging process (5) and is defined by the cessation of cell division and distinctive phenotypic alterations, leading to irreversible proliferation arrest (6, 7). Senescent cells show activation of several characteristic markers, including p53 (cellular tumor antigen p53), p21^Cip1^ (cyclin-dependent kinase inhibitor-1), p16^INK4a^ (cyclin-dependent kinase inhibitor-2A), the increased activity of senescence-associated β-galactosidase (SA-βGal) (8), and the secretion of various bioactive molecules, collectively known as senescence-associated secretory phenotype (SASP) (6). Following chronic exposure to hazardous substances that leads to chronic oxidative stress, COPD lungs show signs of accelerated lung aging with the accumulation of senescent cells, together with the loss of endogenous anti-aging molecules and a persistent state of chronic inflammation that has a similar profile to the SASP (9).

MicroRNAs (miRNAs) are small endogenous non-coding RNAs, between 18-23 nucleotides, that play important roles in gene regulation (10). miRNAs are involved in multiple biological processes, including cellular senescence. Changes in miRNA expression have been described in plasma, sputum, bronchoalveolar lavage fluid and lung tissue of COPD patients (11–13). Specifically, miR-34a and miR-570 levels are elevated in lung tissues and small airway epithelial cells (SAEC) from COPD patients and are strongly implicated in the induction of cellular senescence. MiR-34a and miR-570 are induced by oxidative stress and induce cellular senescence by targeting and suppressing the expression of the anti-aging molecules sirtuin 1 (SIRT1) and SIRT6 (14).

Senescent cells may damage their local environment and induce paracrine senescence in bystander cells due to secretion of the SASP (15). However, the mechanisms for spreading senescence are not fully understood. Extracellular vesicles (EVs) encapsulate specific cargo composed of RNA, miRNA, DNA, proteins or metabolites (16) and have the capacity to trigger molecular and phenotypical changes in recipient cells (17). Previous studies have proposed that EVs can induce cellular senescence in recipient cells, however these studies used mouse cells or transformed cell lines and did not address the mechanism involved (18–21). Modification of EV production and composition has been described in sputum and plasma of COPD subjects, however the function of EVs in the acceleration of lung aging has not been explored (22, 23). Therefore, we hypothesized that EVs transfer cellular senescence by transporting and transferring miR-34a and miR-570 into neighboring cells.

## Materials and Methods

### Cell culture and transfections

Primary small airway epithelial cells (SAEC) were cultured as monolayers in LHC-9 medium (Invitrogen) on collagen (1% w/v) coated plates. Cells were extracted from lung tissue from subjects undergoing lung resection surgery at the Royal Brompton Hospital, London. The subjects were matched for age and smokers and COPD subjects for smoking history (**Online Data Supplement Table 1**). All subjects gave informed written consent, and the study was approved by the London-Chelsea Research Ethics committee (study 15/SC/0101). Recipient SAEC were transfected with mirVana miRNA inhibitor (miRVanaTM miRNA inhibitor Negative control #1, hsa-miR-34a MH11030) (Ambion, Life Technologies, Foster City, CA) using Lipofectamine RNAimax.

### Isolation of extracellular vesicles

Conditioned media from BEAS2B cells or SAEC was centrifuged at 300*g* for 10 min to eliminate debris, then centrifuged at 20000*g* for 30min to isolate large EVs and at 100000*g* for 2h for small EVs. The resultant pellets were resuspended in phosphate buffered saline (PBS) or media. The expression of markers of EVs, CD9, CD81, CD63, Alix and calreticulin were analyzed by Western Blot. EV concentration was measured by NanoFCM (NanoFCM, Xiamen, China) and by flow cytometry with DynabeadsTM coated with a primary monoclonal antibody specific for CD9 (Invitrogen, Waltham, MA) (**Online Data Supplement Material and Methods**). Large and small EVs treated for 20 min with 0.01% v/v Triton-X100 were used as an internal control.

### Stimulation of recipient cells with EVs

EVs isolated from SAEC were resuspended in 300μl of media. Large EVs concentration varied from 1.86×10^8^ and 2.66×10^11^ and small EVs between 9.30×10^8^ and 1.46×10^13^. To closely reflect normal physiology, the volume of EVs was normalized to the number of donor cells, and 150μl of EVs-solution (∼2 x 10^11^ particles) was applied to 250 000 recipient SAEC plated in 6-well plates. The expression of miR-34a was analyzed after 3h, and the expression of SIRT1 and p21^CIP1^ were analyzed after 12h by qPCR analysis. The protein expression of SIRT1 and p21^CIP1^ was analyzed after 48h by Western Blot. The activity of SA-β-galactosidase was analyzed by colorimetric analysis after 48h (**Online Data Supplement Material and Methods**).

### Analysis of cell uptake of EVs

EVs were stained with the fluorescent lipid membrane label, PKH67, according to the manufacturer’s instruction (Sigma Aldrich, St Louis, MO). The washes were recovered and used as a control for dye leakage. BEAS2B cells and SAEC were stimulated with 100μl of EVs stained solution for 3, 12, 24 or 48 h at 37°C with 5% (v/v) CO_2_. Cells were fixed and stained with Red Cell trace (Life technologies, Foster City, CA) and DAPI (Abcam, Cambridge, United Kingdom) and imaged by confocal microscopy. Cells were trypsinized and PKH67 fluorescence was detected into BEAS2B cells on a Canto II flow cytometer (Beckton Dickinson Biosciences, Franklin Lakes, NJ).

### Statistical analysis

Data are expressed as mean ± SEM. Results were analyzed using two-way ANOVA with Bonferroni post-test correction, Mann-Whitney, Wilcoxon or Kruskal-Wallis tests as appropriate. GraphPad Prism 9.2 software (GraphPad software, La Jolla, CA) was used for analysis. Values of p≤0.05 were considered statistically significant.

## Results

### SAEC from COPD subjects produce EVs enriched with miR-34a

Large and small EVs were isolated from cell media by ultracentrifugation and their concentration was assessed by Nanoflow cytometry. EVs were characterized by western blot and the vesicular structure of EVs was confirmed by flow cytometry (**Online Data Supplement Figure 1**). SAEC from COPD subjects produced 10-fold more large vesicles than EVs derived from healthy SAEC (**Figure 1A**) (healthy: 1.13 ± 0.35×10^10^, n=5 *vs* COPD: 1.35 ± 0.99×10^11^ particles/ml, n=5), with no difference in their average size (**Figure 1B**). Similarly, COPD SAEC produced more small EVs than SAEC from healthy donors (**Figure 1C**) (healthy: 2.2 ± 0.21×10^12^, n=5 *vs*. COPD: 1.37 ± 0.39×10^12^ particles/ml, n=5) but not significantly (p=0.055) and again there was no difference in their size (**Figure 1D**). Moreover, SAEC from COPD subjects expressed more miR-34a compared to SAEC from healthy donors (**Figure 1E**) and this was also reflected in EVs with both large and small EVs derived from COPD SAEC showing enrichment for miR-34a compared to EVs derived from healthy cells (**Figure 1F-G**). The expression of miR-570 was increased in COPD SAEC compared to non-smoker cells (**Online Data Supplement Figure 2A**), but no difference was detected in large or small EV (**Online Data Supplement Figure 2B and 2C**). As these differences between healthy and COPD SAEC were detected at baseline, we examined whether oxidative stress could trigger production of EVs using the airway epithelial cell line BEAS2B incubated with H_2_O_2._ In response to oxidative stress, the release of the large and small EVs was not modified with no modification of their average size (**Online Data Supplement Figure 3A-D**). As previously shown, expression of miR-34a was significantly increased in BEAS-2B cells after oxidative stress (**Online Data Supplement Figure 3E**) (14), and despite numbers of EVs not increasing, both large and small EVs were significantly enriched with miR-34a following H_2_O_2_ exposure (**Online Data Supplement Figures 3F and 3G**). Similarly, expression of miR-570 was also significantly increased in BEAS2B cells stimulated with H_2_O_2_ (**Online Data Supplement Figure 3H)**, but this cellular change was not reflected in either large or small EVs derived from these cells (**Online Data Supplement Figures 3I and 3J**). Thus, oxidative stress alone does not seem to trigger the release of EVs from airway epithelial cells but may stimulate the expression and the packaging of miR-34a within EVs.

**Figure 1:**
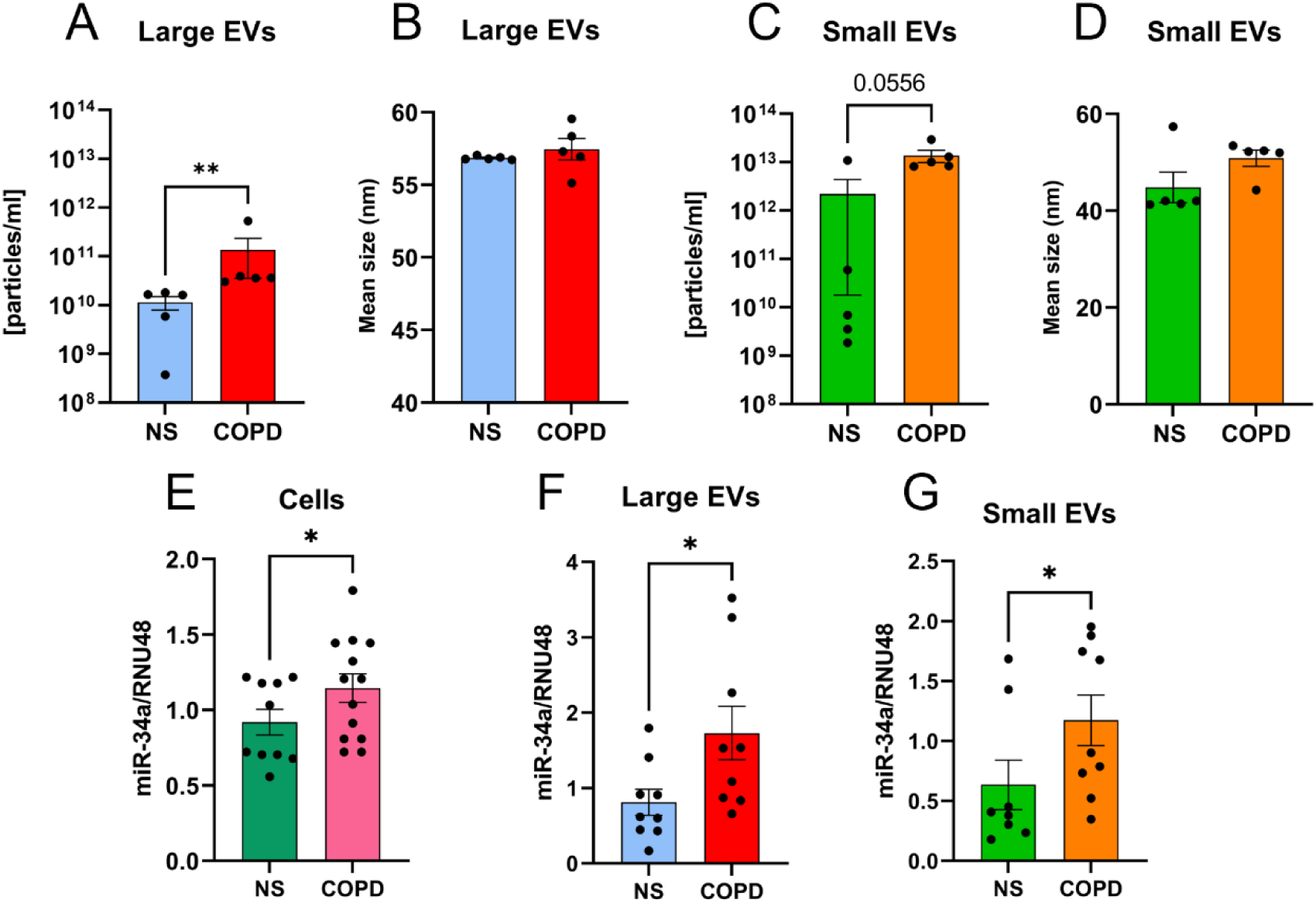
COPD SAEC produce EVs enriched with miR-34a. Large and small EVs produced by healthy and COPD SAEC were isolated from 7-days conditioned media. The concentration of EVs (**A, C**), and size (**B, D**) were analyzed by NanoFCM. Total RNA was extracted and miR-34a expression was determined in cells (**E**, n=12), large EVs (**F**, n=9) and small EVs (**G**, n=9) by qRT-PCR. Each point represents a different patient. Data are mean ± SEM, analyzed by Mann-Whitney test. *p<0.05, **<0.01

### EVs produced by SAEC are taken up by healthy recipient SAEC

To determine whether EVs can be taken up by epithelial cells, EVs were isolated from SAEC cultured for 7-days and labelled with the fluorescent dye PKH67. Recipient healthy SAEC cells were treated with PKH67-labelled EVs for up to 16h and analyzed by flow cytometry. Internalization of EVs was time-dependent as after 3h and 16h exposure to large EVs, the proportion of recipient SAEC positive for PKH67 compared to untreated cells increased from 5.7 ± 0.59% to 53.6 ± 0.83% (**Figure 2A**), with no difference between the EVs derived from COPD- and healthy-SAEC. A similar pattern was seen for the uptake of small EVs (3h: 38.7 ± 10.2% positive cells; 16h: 92.7 ± 0.08% positive cells), where the proportion of recipient cells containing EVs being was greater than that for larger EVs (**Figure 2B**). The proportion of recipient cells positive for PKH67 was not different when exposed to small EVs-derived from COPD SAEC compared to healthy-derived EVs (**Figure 2B**). Uptake is an active process as EVs were not detectable in recipient cells when incubated at 4°C. Internalization was confirmed following exposure of cells to PKH67-labelled EVs for 6h using fluorescence microscopy (**Figure 2C**) and was lost when EVs were treated with the detergent Triton X100 (**Figure 2D**). These data suggest that, within 3h of incubation, EVs are taken up by healthy recipient SAEC.

**Figure 2:**
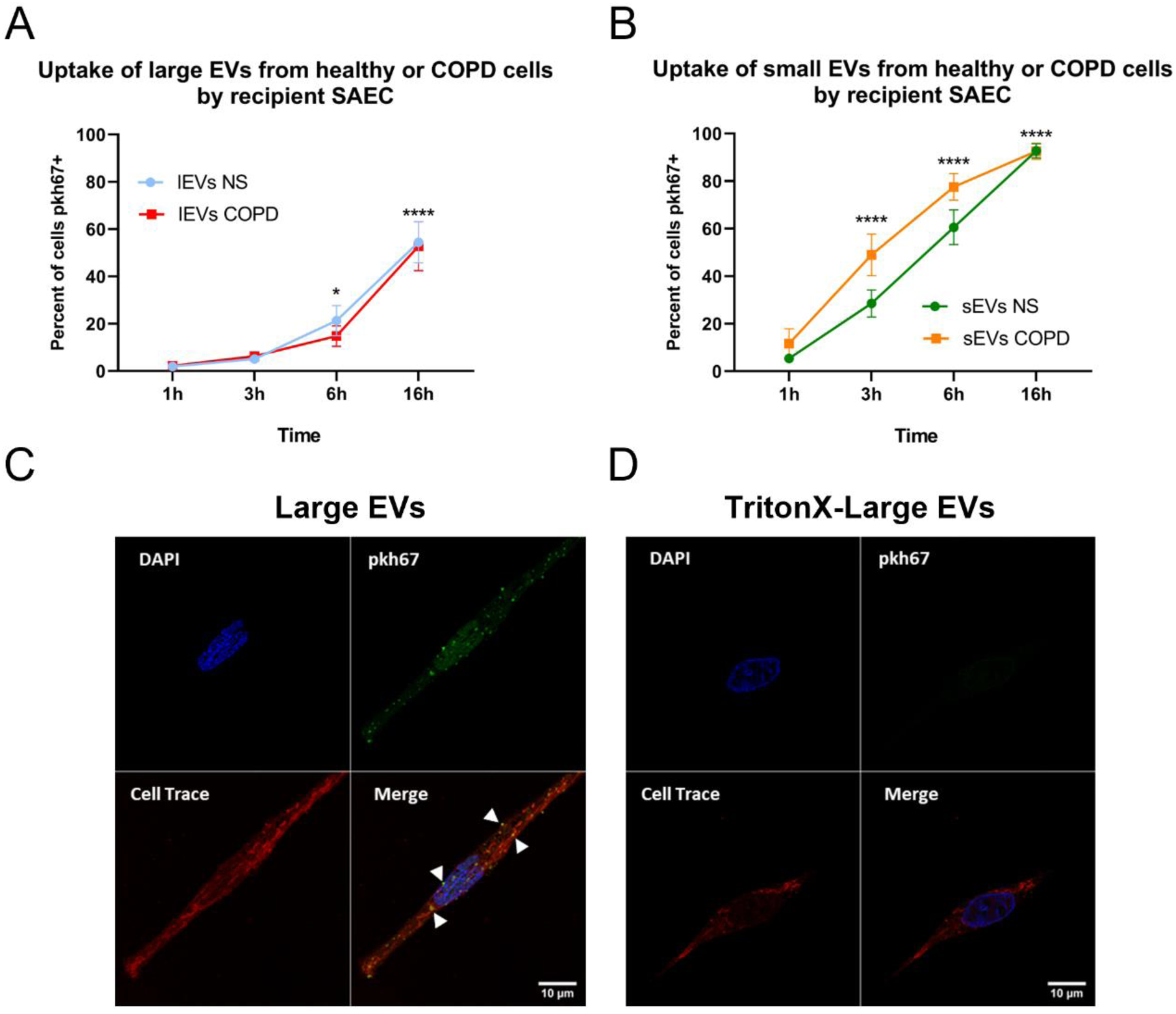
Extracellular vesicles are uptake by recipient SAEC. Large and small EVs were isolated from healthy or COPD SAEC media, stained with PKH67 and resuspended in 300μl of media. Healthy SAEC were stimulated with PKH67-labelled large (**A**) or small (**B**) EVs and the uptake of EVs was analyzed by flow cytometry at 1h, 3h, 6h and 16h. PKH67-labelled large EVs (**C**) or PKH67-labelled EVs treated with TritonX100 (**D**) were incubated with 125 000 recipient healthy SAEC for 6h. Cells were fixed and nuclei stained with DAPI (blue) and cytoplasm with cell tracker (red) and imaged by fluorescence microscopy. Images are representative of 4 experiments. Data are expressed as mean ± SEM, analyzed by Two-way ANOVA with post hoc Sidak *p<0.05, ****p<0.0001

### COPD SAEC-derived EVs transfer functional miRNA-34a into healthy SAEC

We next examined the functional effect of EVs on recipient cells by incubating 2.5 x 10^5^ healthy SAEC with 150μl of EVs (∼2 x10^11^ vesicles) derived from either healthy or COPD SAEC (**Figure 3A**). Transfer of large, but not small EVs from COPD SAEC significantly increased the expression of miR-34a (**Figure 3B and Online Data Supplement Figure 4A**) and decreased the expression of SIRT1 mRNA (**Figure 3C and Online Data Supplement Figure 4B**) in recipient cells compared to EVs derived from healthy SAEC. This effect was due to large EVs as there was no modification in miR-34a and SIRT1 mRNA expression in recipient cells treated with EVs previously exposed to Triton X-100 or with media that had been depleted of EVs (**Figures 3D and 3E**). In the same way, transfer of large, but not small EVs diminished the protein level of SIRT1 in recipient SAEC and this reduction in SIRT1 was lost when EVs were dissolved by Triton or with media depleted of EVs (**Figure 3F and Online Data Supplement Figure 4C**). It remained a possibility that the presence of large EVs could induce endogenous production of miR-34a in recipient cells. To address this, recipient cells were pre-treated with transcriptional inhibitor actinomycin D, prior to exposure to EVs (**Figure 3G**). This did not affect the increased miR-34a expression observed in response to EVs from COPD subjects and supported the concept that miR-34a is transferred from EVs and does not result from endogenous synthesis by recipient cells.

**Figure 3:**
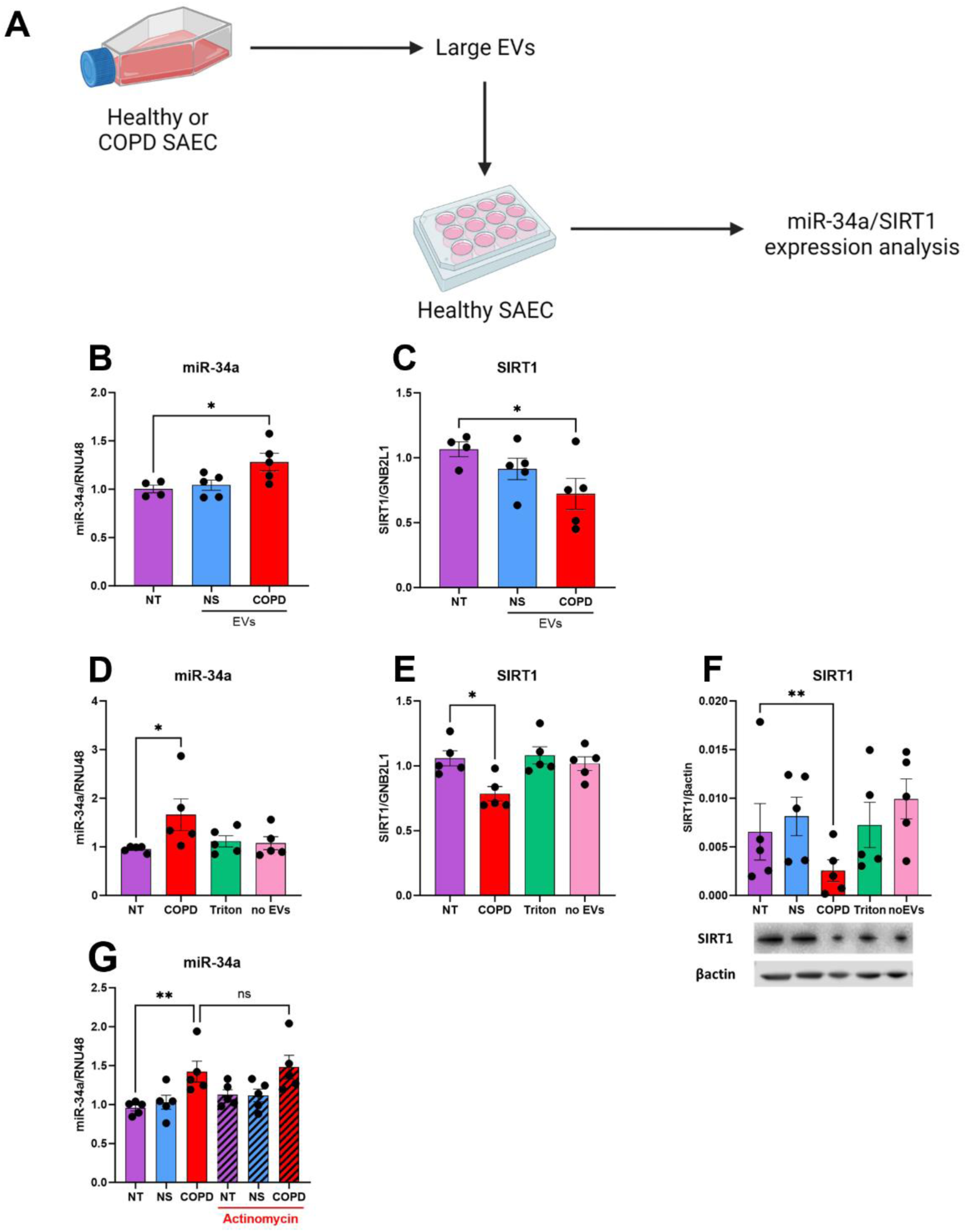
Large EVs from COPD SAEC transfer functional miR-34a in recipient small airway epithelial cells. Large EVs were isolated from 7-days conditioned media from COPD or non-smoker SAEC. (**A**) 250 000 recipient healthy SAEC were stimulated with 150μl of large EVs (∼2 x 1011 particles), with EVs treated with Triton X100 (Triton), or with media where EVs have been depleted (SN-EVs). Expression of miR-34a (**B, E**), and SIRT1 (**C, F**) were measured in recipient cells after 3h (for miR-34a) and 12h (SIRT1) by RT-qPCR. After 48h stimulation, (**D**) SIRT1 protein was analyzed by western blotting. (**G**) Recipient BEAS2B cells were pre-treated with 10μM of actinomycin D for 3h before being treated with EVs from NS or COPD subjects. miR-34a expression was measured after 3h by RT-qPCR. Data are expressed as mean ± SEM, analyzed by Kruskal-Wallis test with post hoc Dunn’s or Mann-Whitney as appropriate where *p<0.05 ** p<0.01

Taken together, these data suggest that large EVs produced by COPD SAEC transfer miR-34a into recipient epithelial cells and have the capacity to induce cellular senescence by inhibiting SIRT1.

### COPD SAEC-derived EVs induce senescence markers into healthy SAEC

To assess whether EVs-mediated reduction in SIRT1 led to an expected increase in markers of senescence, expression of p21^CIP1^, IL-6 and the activity of SA-β-gal were measured in recipient cells following incubation with EVs from COPD SAEC. Transfer of large EVs from COPD subjects significantly increased mRNA and protein expression of p21^CIP1^ in recipient cells (**Figure 4A and 4B**). Moreover, recipient cells treated with COPD large EVs produce significantly more IL-6 than untreated cells (**Figure 4C**). Recipient cells treated with large EVs from COPD SAEC also exhibit increased SA-β-galactosidase (SA-βGal) compared to untreated cells, which was lost when EVs were treated with Triton X100 (**Figure 4D**).

**Figure 4:**
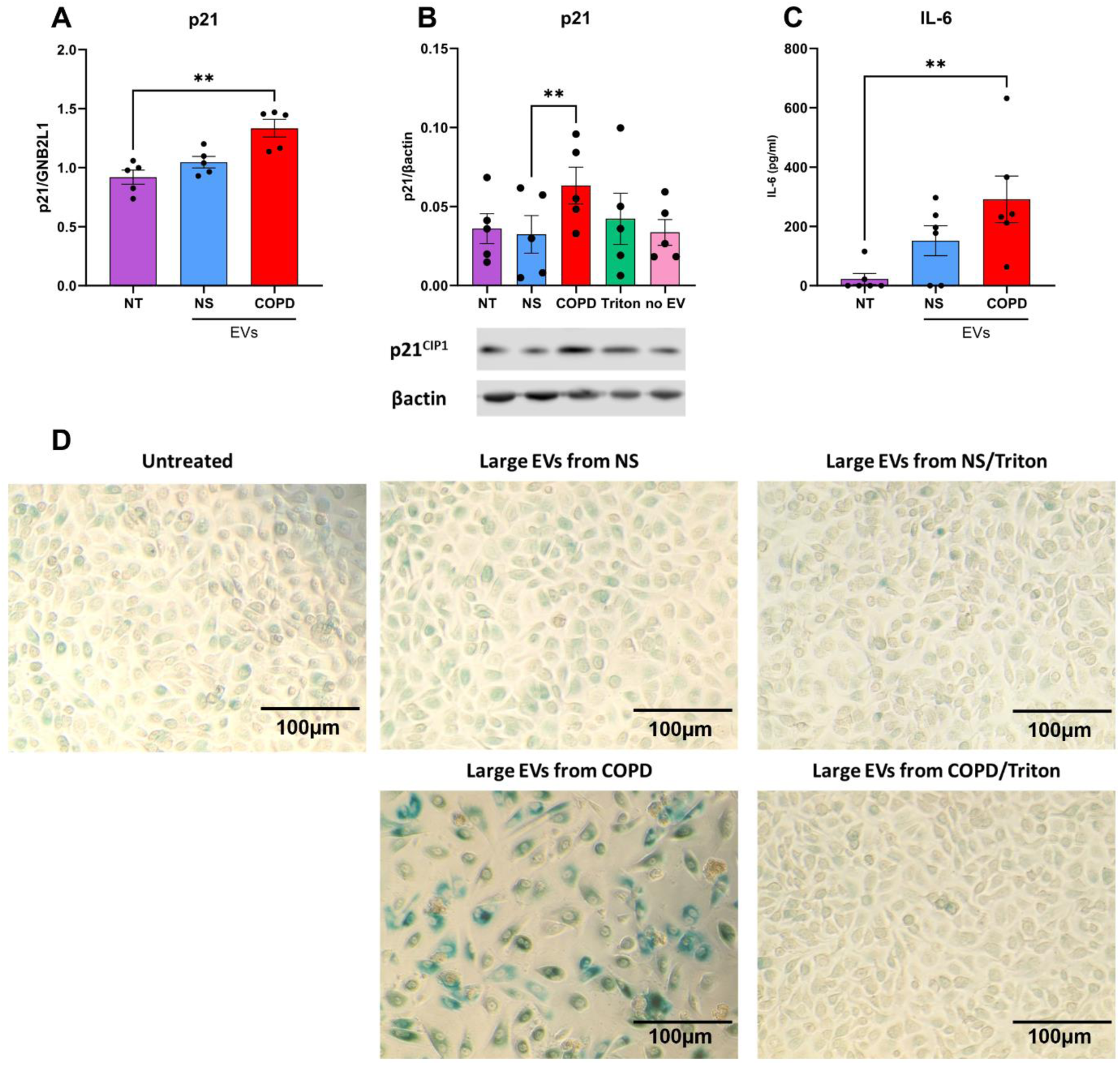
Large EVs from COPD SAEC induce senescence markers in recipient small airway epithelial cells. EVs were isolated from 7-days conditioned media from COPD or non-smoker SAEC. 250 000 recipient healthy SAEC were stimulated with 150μl of large EVs (∼2 x 1011 particles) from NS or COPD donors. Expression of p21 was measured in recipient cells by (**A**) RT-qPCR after 12h and by (**B**) western blotting after 48h. (**C**) IL-6 secretion was measured in recipient cell media by ELISA after 48h. (**D**) Recipient SAEC were stained for senescence associated β-galactosidase and imaged by light microscopy. Images are representative of 4 experiments. Each point represents EVs isolated from different SAEC donors (n=6). Data are expressed as mean ± SEM, analyzed by Kruskal-Wallis test with post hoc Dunn’s or Mann-Whitney as appropriate where *p<0.05 ** p<0.01

p21^CIP1^ mRNA expression was also significantly increased in recipient cells exposed to COPD small EVs compared to non-smoker small EVs (**Online Data Supplement Figure 5A**), but no effect of COPD small EVs was detected on protein expression of p21^CIP1^ in recipient cells after 48h (**Online Data Supplement Figure 5B**). Similarly, IL-6 production was not modified in recipient cells treated with small EVs (**Online Data Supplement Figure 5C**).

Taken together, these data suggest that large EVs produced by COPD SAEC induce the expression of p21^CIP1^, production of the SASP factor IL-6, and increased activity of SA-βGal confirming the potential role of EVs in the induction of cellular senescence.

### COPD SAEC-derived EVs induce senescence markers in recipient SAEC through miR-34a transfer

Having identified EVs as a potential vector of cellular senescence, we aimed to confirm the involvement of miR-34a pathway. SAEC from healthy donors were transfected overnight with an antagomir against miR-34a or a control scrambled sequence before being incubated with large EVs derived from healthy or COPD SAEC (**Figure 5A**). After 3h exposure to EVs, the expression of miR-34a induced by large EVs from COPD cells was significantly abrogated in recipient cells previously transfected with the antagomir compared to cells transfected with a control sequence (**Figure 5B**). Consequently, recipient cell transfection with the antagomir prevented reduction of SIRT1 mRNA and protein expression (**Figure 5C and 5D**) and blocked the augmentation of p21^CIP1^ mRNA and protein expression in recipient SAEC (**Figure 5E and 5F**). To further confirm the role of miR-34a contained in EVs in the induction of cellular senescence, recipient cells were treated with EVs derived from donor healthy and COPD cells transfected with an antagomir against miR-34a (**Online Data Supplement Figure 6A**). Large EVs from control COPD SAEC increased miR-34a expression in the recipient cells which is blocked when donor cells are transfected with the antagomir (**Online Data Supplement Figure 6B**). Consequently, treatment of recipient cells treated with EVs isolated from COPD SAEC-antimiR34a prevented reduction of SIRT1 mRNA and augmentation of p21^CIP1^ mRNA expression (**Online Data Supplement Figure 6C and 6D**). Taken together, these data suggest that transfer of miR-34a via large EVs produced by COPD SAEC induces cellular senescence in recipient healthy cells by inhibiting the expression of SIRT1 and subsequently increasing expression of p21^CIP1^. Furthermore, pre-treatment of SAEC with an antagomir against miR-34a is sufficient to inhibit these effects.

**Figure 5:**
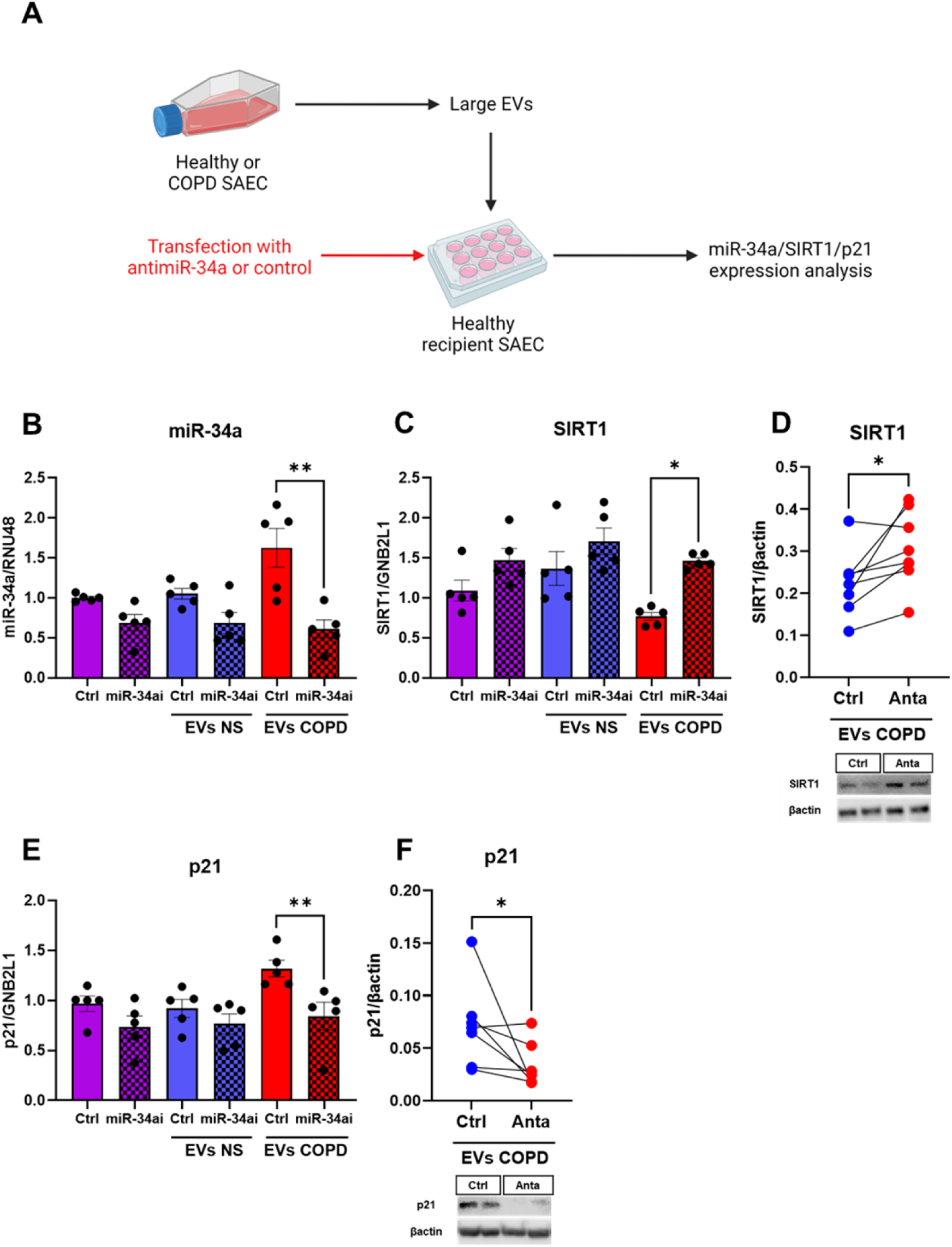
Large EVs from COPD SAEC induce senescence through miR-34a transfer. (**A**) Recipient healthy SAEC were transfected with an antagomir against miR-34a or with a control sequence overnight prior to treatment with large EVs isolated from NS or COPD SAEC cell media. Changes in (**B**) miR-34a, (**C**) SIRT1 and (**E**) p21 mRNA were measured by qRT-PCR after 3 and 12 h stimulation. (**D**) SIRT1 and (**F**) p21 protein expression were analyzed 48 h after recipient SAEC stimulation by western blotting. The band density of each blot is represented as histogram. Each point represents treatment with EVs isolated from different SAEC donors. Data are expressed as mean ± SEM, analyzed by Wilcoxon test. *p<0.05

## Discussion

COPD is associated with accelerated aging. Normal and pathological (accelerated or premature) aging leads to the accumulation of comorbidities and the consequent deterioration of health and quality of life (24). A major hallmark of aging is cellular senescence (25). Recent studies propose that senescent cells are capable of inducing neighboring cells to undergo senescence through cell-to-cell contact or in a paracrine fashion (26, 27). The possible role of EVs in the induction of paracrine senescence is suggested by some studies, however the mechanism has not been fully determined (28, 29). Our experimental results provide a proof of concept that EVs produced by SAEC transfer miR-34a and induce cellular senescence in recipient epithelial cells. Thereby EVs and associated miR-34a may contribute to the spread of senescence and hence aging. As such this mechanism may contribute to the pathophysiology of COPD, but more broadly to age-related diseases and to the development of multimorbidity in the elderly.

The function of EVs in aging and in COPD is currently uncertain. Contradictory results have been described depending on the methods used to isolate EVs (30, 31), confirming that the analysis methods used to characterize and describe EVs are of utmost importance. Ultracentrifugation is the most commonly used method for EVs purification (32), however it can lead to the sedimentation of large proteins and/or proteins that are non-specifically associated with EVs. Other methods, such as size exclusion chromatography, can be used to isolate different populations of EVs but may also lead to deformation and rupture of EVs that can be accompanied by a significant decrease on the overall EV number (32, 33). Thus, and in accordance with current guidelines, we have tested several speeds and durations of ultracentrifugation and carefully selected the method that allowed the isolation of EVs (34). We have confirmed the lipid nature of EVs by analyzing the expression of specific markers and using a detergent to disrupt the vesicles (**Online Data Supplement Figure 1**). We have then shown that SAEC from COPD subject produce more EVs than SAEC from age-matched healthy subjects based on nanoflow cytometry (**Figure 1A-D**). In agreement with these results, changes in EV production have been demonstrated in age-related diseases. The concentration of CD9 positive EVs is significantly elevated in the plasma of COPD subjects and is correlated with pro-inflammatory factors including C-reactive protein (CRP) and IL-6 (35).

The composition of EVs may also be altered in COPD and in aging, possibly due to oxidative damage of donor cells. Proteomic analysis of EVs produced by BEAS2B revealed that 33% of proteins were differentially expressed after exposure to cigarette smoke extract, with up-regulation of proteins involved in cell-to-cell communication and immune responses (36). Moreover, stimulation of BEAS2B cells *in vitro* with cigarette smoke extract leads to the production of EVs containing miR-210 (37). The comparison of miRNA expression in EVs isolated from bronchoalveolar lavage fluid obtained from non-smokers and smokers showed that smoking alters the miRNA profile, with increased the expression of miR-21 and miR-27a (38). In agreement with previous publications, we have shown that treatment of BEAS2B cells with hydrogen peroxide induces miR-34a expression in cells and EVs. However, without any additional stimulation, EVs derived from COPD SAEC are enriched with miR-34a. Once the disease is established, oxidative stress persists in cells even after smoking cessation due to persistent inflammation, or impaired endogenous antioxidant defenses (39). Thus, oxidative stress may be essential in the expression and packaging of miR-34a in EVs.

Vesicular miRNAs are mediators of genetic exchange between cells. Following delivery, miRNAs regulate the translation of target genes and ultimately the function of the recipient cells (40). Our data showed that miR-34a associated with EVs can be transferred into recipient SAEC to suppress expression of SIRT1, leading to increased expression of senescence markers. Our results are supported by a study showing that EVs produced by mouse myoblasts in response to oxidative stress contain miR-34a that can be transferred into bone marrow mesenchymal (stromal) cells with an increase in the senescence biomarker SA-βGal (18). MiR-34a is ubiquitously expressed and is involved in numerous cellular functions, including proliferation, epithelial to mesenchymal transition, apoptosis, migration, cell cycle and cellular senescence (41) and is thus implicated in numerous age-related diseases, including cancer (41), neurodegenerative diseases (42), and cardiovascular diseases (44). Although the composition of EVs reflects the pathophysiological state of the source cell, miRNA content is often markedly different from the miRNA content of the parent cells (45, 46). These findings are consistent with our data showing an increased expression of miR-34a and miR-570 in source SAEC from COPD subjects but only an increased expression of miR-34a in EVs derived from COPD cells. It implies that miRNAs are not passively sorted in EVs but are selectively and actively packaged. Altogether, these data confirm that vesicular miRNAs, and particularly, miR-34a is an important modulator of cellular senescence that may actively participate in the development of age-related diseases. Thus, measure of vesicular miR-34a in bronchoalveolar lavage, sputum or plasma may be a novel biomarker of accelerated aging.

Previously, the spread of cellular senescence has been ascribed to components of the SASP, which *in vitro* may induce senescence in healthy cells (27, 47). However, our study does not support this hypothesis as EVs-depleted cell media and the disruption of vesicles with a detergent failed to reduce SIRT1 or induce an increase in p21^CIP1^ in recipient cells. This suggests that EVs are a more effective means of propagating senescence in small airway epithelial cells.

In conclusion, our data demonstrate that EVs act as vehicles that are central in the induction of cellular senescence and participate in the development of COPD and age-related diseases. We showed that miR-34a could be packed into EVs produced by SAEC and be actively transferred into healthy SAEC consequently promoting the downregulation of SIRT1 and an increase in cellular senescence. An elegant study from Tian et *al* revealed that aging is a multisystem process whereby the aging of one organ can influence the aging of multiple other systems. Notably, they have shown that the aging of the pulmonary systems leads to faster cardiovascular aging, which in turn results in faster aging of the musculoskeletal and renal system (48). Thus, by spreading senescence locally, EVs may participate in COPD progression, but may also spread senescence to other organs and account for comorbidities and multimorbidity of the elderly. As EVs are detectable in every body fluid and have a longer half-life than free RNA, they can be measured in biological fluids as a biomarker of diseases of accelerated aging and of multimorbidity. EVs may also be utilized as a delivery system for miRNA or antagomirs and could be engineered to target specific types of cell. Thus, targeting vesicular miR-34a to reverse cellular senescence could be a novel therapeutic target in the treatment of COPD and age-related diseases.

## Acknowledgment

The authors thank Prof Maria Belvisi (National Heart and Lung Institute, London, UK) and Dr Ken Grime (AstraZeneca, Cambridge, UK). This work was supported by AstraZeneca. The authors acknowledge the NHLI Facility for Imaging by Light Microscopy (FILM) and the Flow Cytometry Facility for use of their confocal microscope and Nano Flow cytometer. This study was supported by the Royal Brompton and Harefield NHS Foundation Trust and Imperial College London.

## Source of grants

AstraZeneca

---

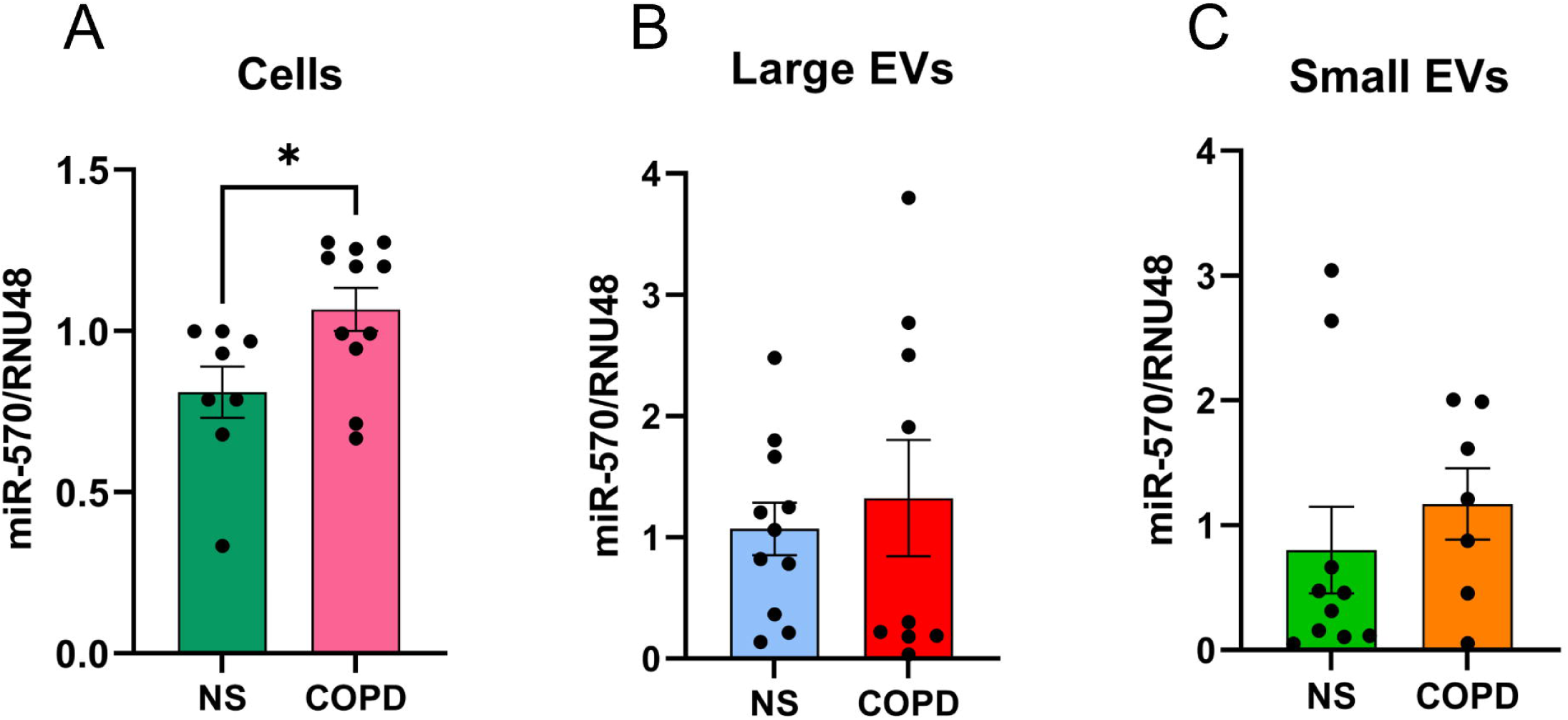

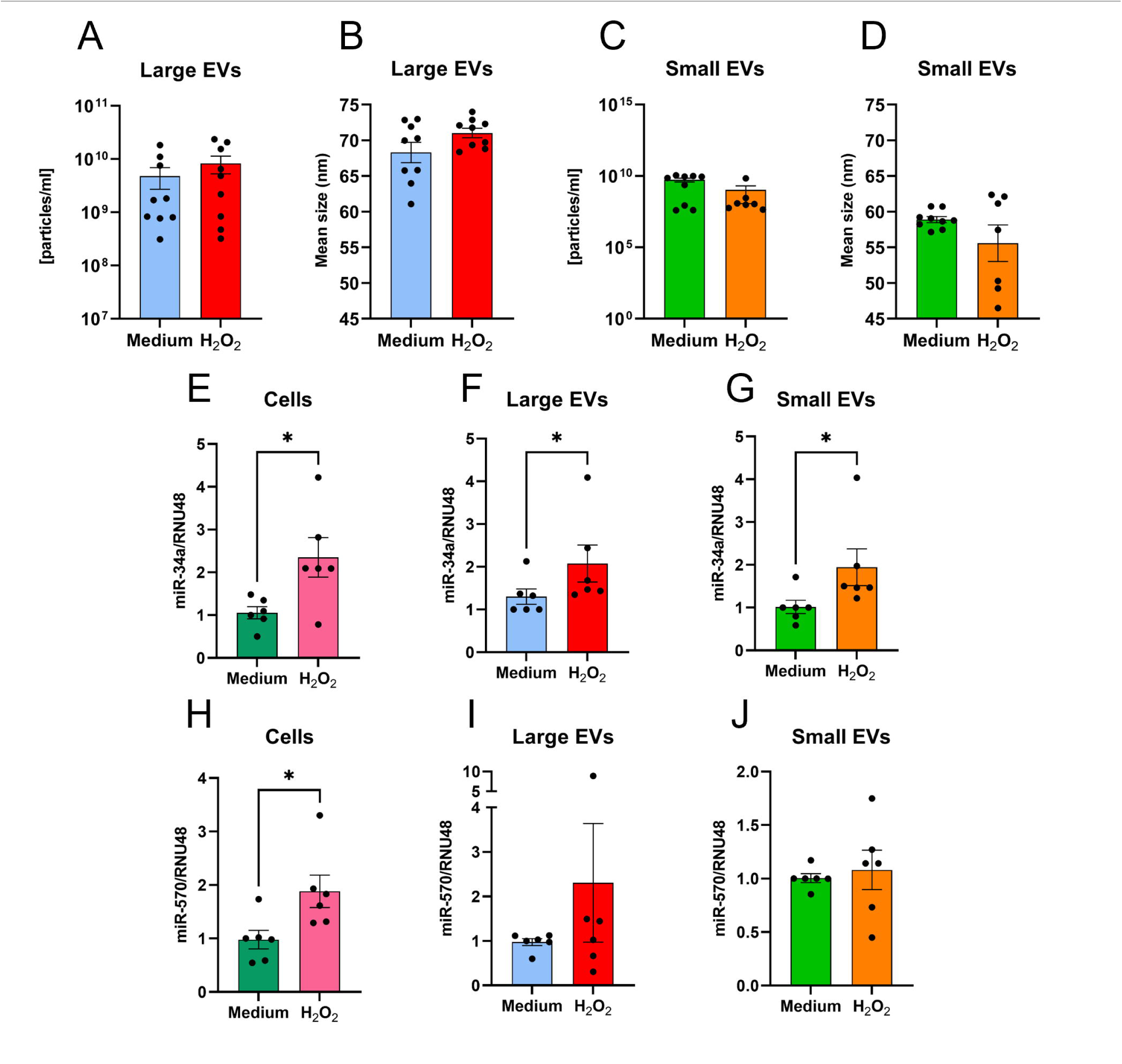

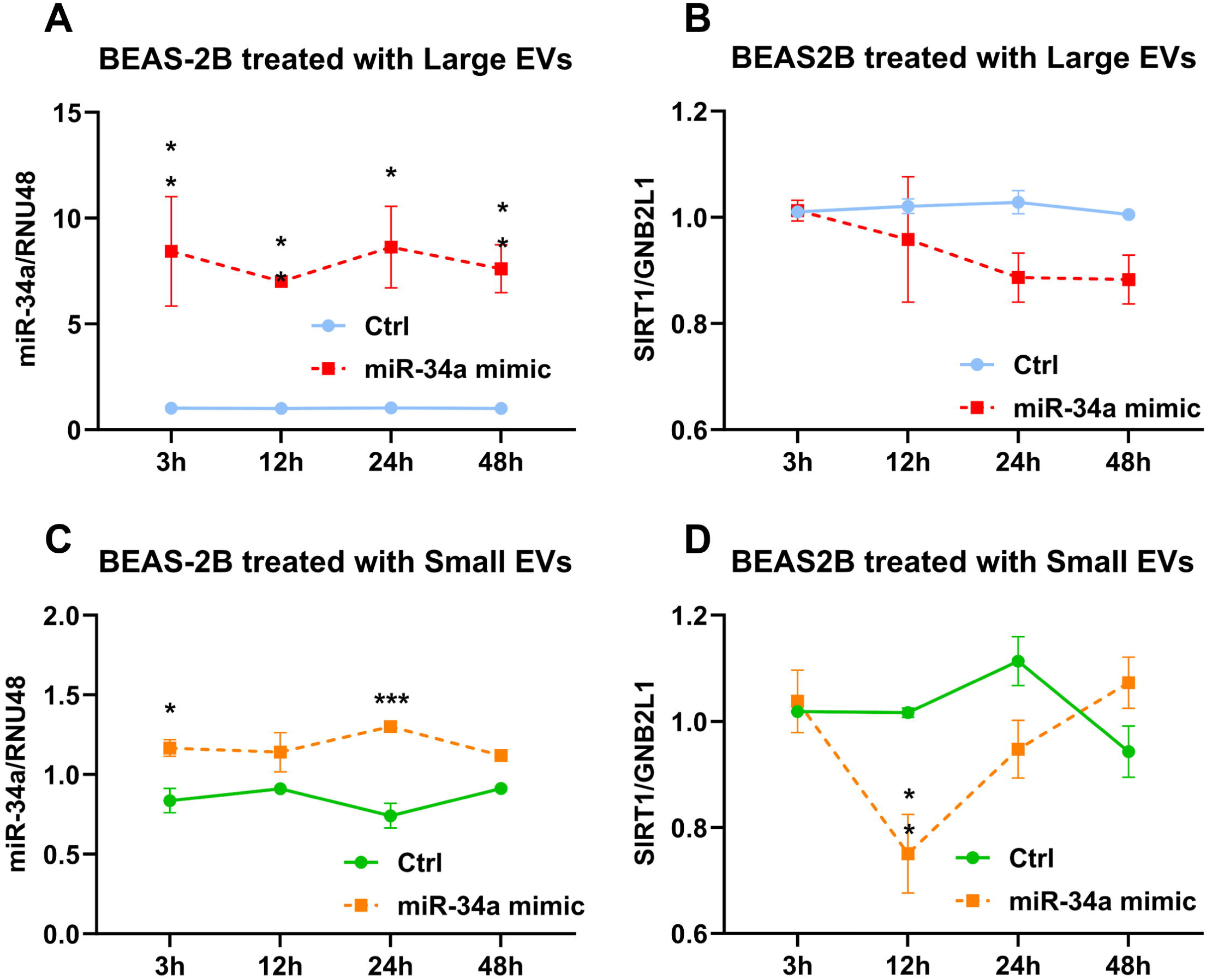

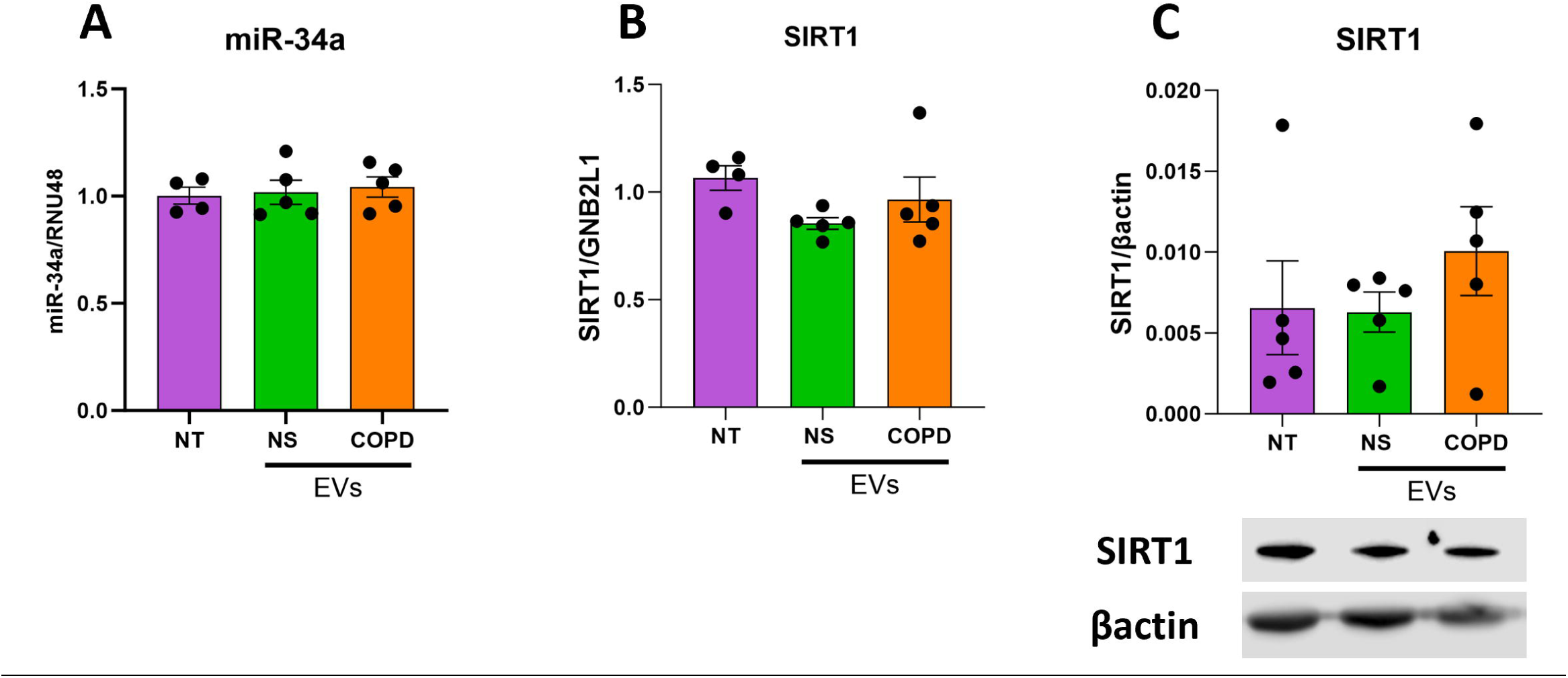

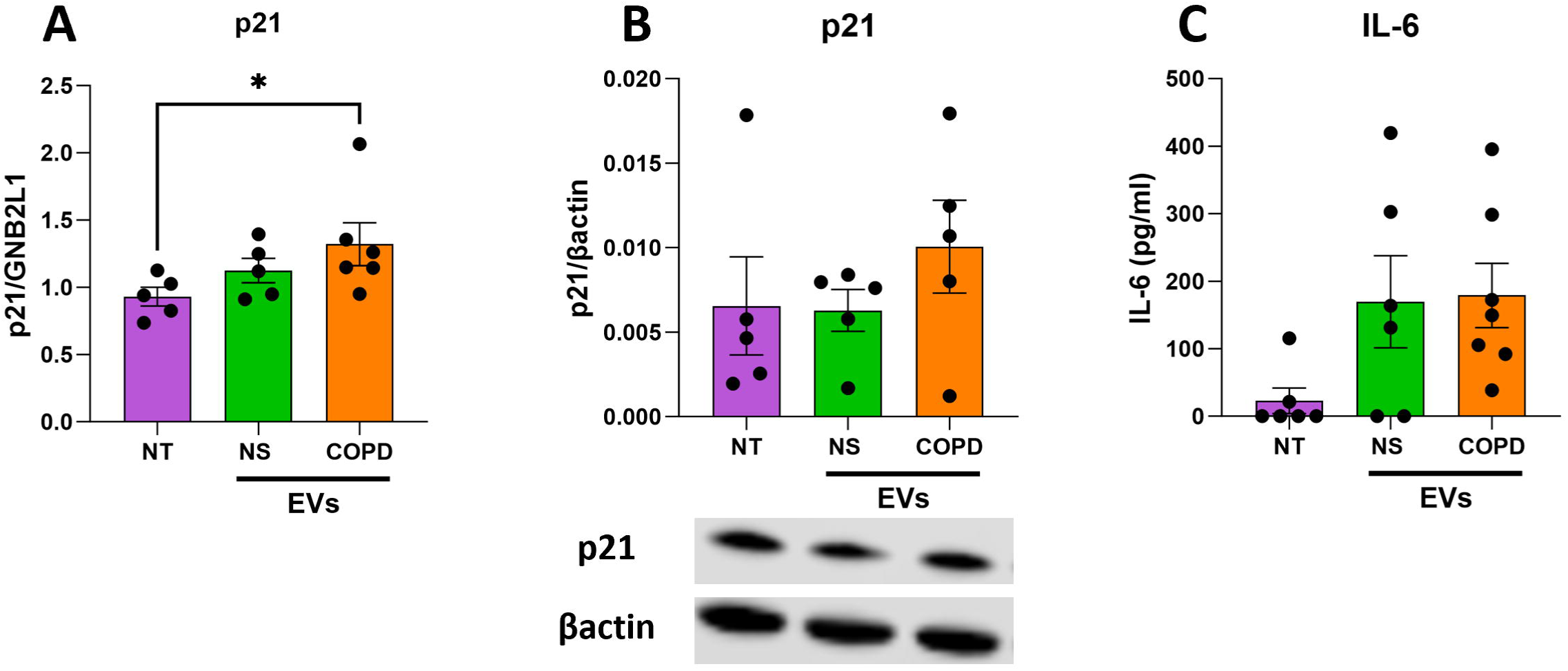

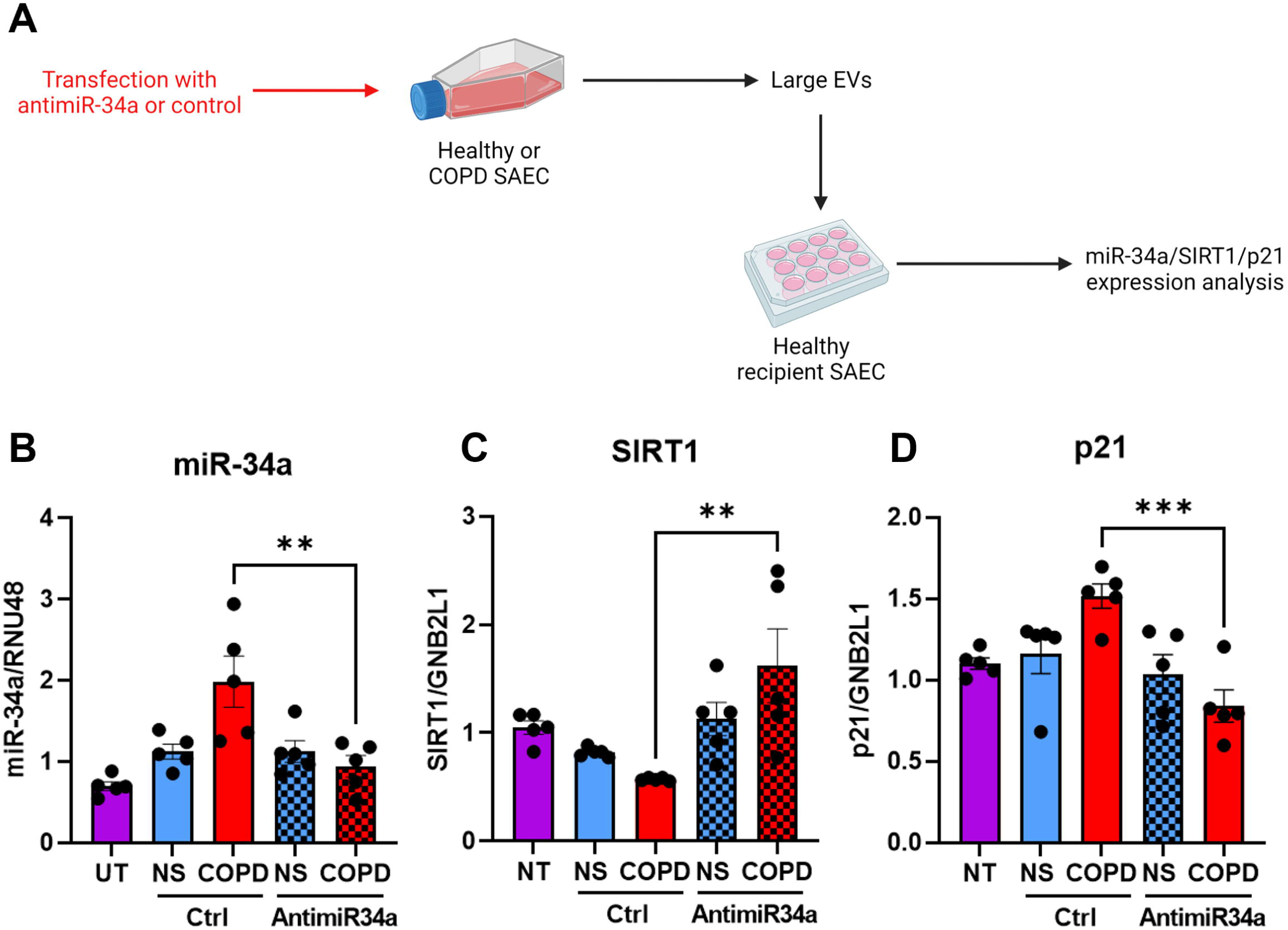

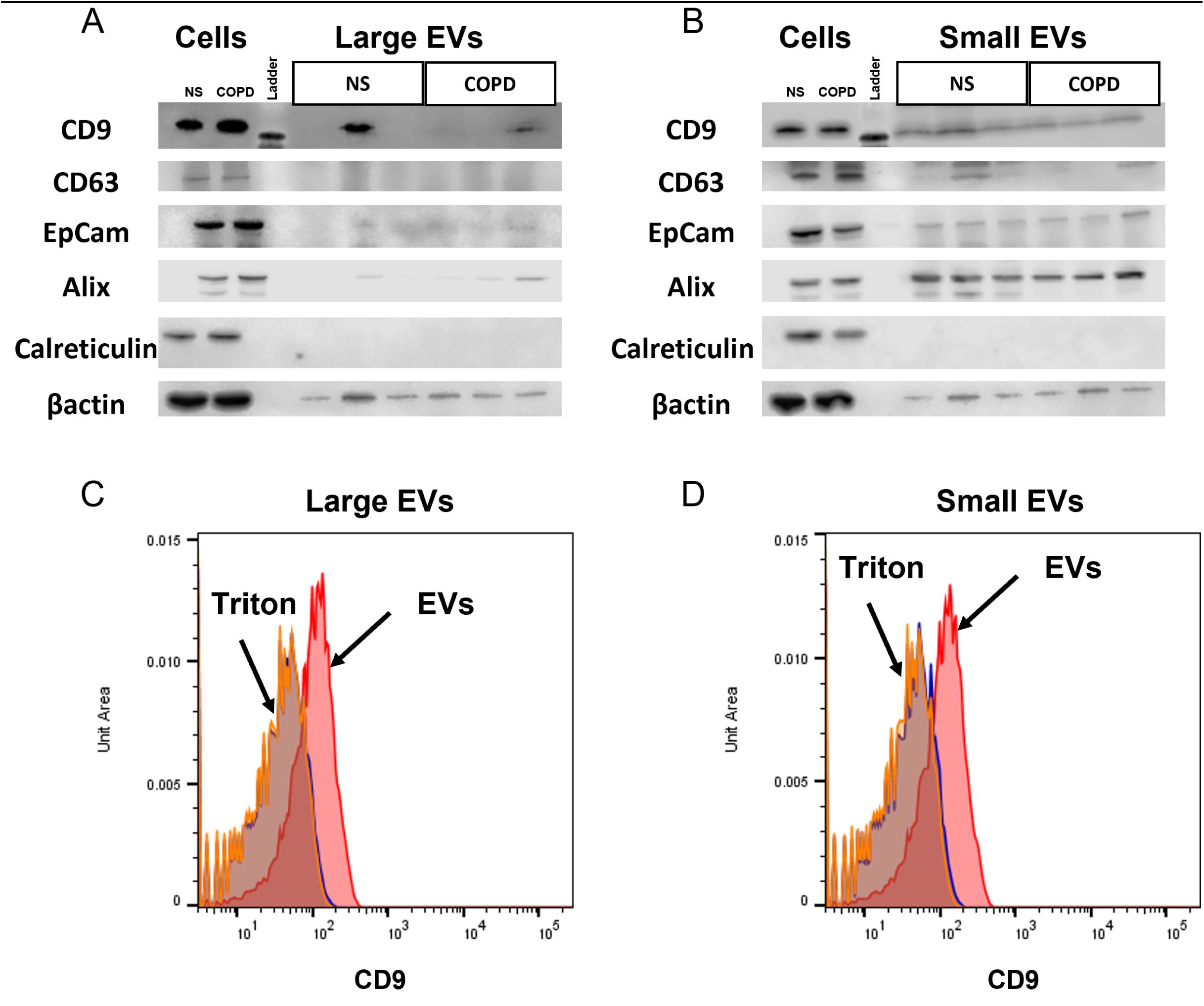

## Notes

### Competing Interest Statement

The authors have declared no competing interest.

