## Supplementary information for "Extracellular vesicles propagate aging in COPD airway epithelial cells by transfer of microRNA-34a"

### Supplemental Table 1

| Characteristics | Non-smoker (n=11) | COPD (n=11) |
|---|---|---|
| Age (years) | 68.6 ± 12.8 | 62.3 ± 7.4 |
| Sex (M:F) | 03:08 | 05:06 |
| $FEV_1$ (L) | 2.1 ± 0.5 | 1.1 ± 0.7 |
| $FEV_1$ (% predicted) | 93.1 ± 14.3 | 37.8 ± 25.3 |
| FVC (L) | 2.8 ± 0.7 | 3.1 ± 1.1 |
| $FEV_1$:FVC | 0.78 ± 0.08 | 0.37 ± 0.14 |
| Pack-years[a] | 0 | 31.91 ± 20.72 |

**Characteristics of study subjects**.

Patients with COPD were categorised according to Global Initiative for Chronic Obstructive Lung Diseases. Definitions. M, male; F, female; $FEV_1$, forced expiratory volume in 1s; FVC, forced vital capacity; [a]Number of cigarettes smoked per day/20 x duration of smoking. Data are expressed as mean ± SEM.

**Supplemental Material and Methods**

### Reagents and antibodies

Hydrogen peroxide ($H_2O_2$) was purchased from Sigma (Poole, UK). Antibodies against the following were used for immunoblotting: βactin (Santa Cruz Biotechnology, Santa Cruz, CA), sirtuin-1 (1F3, MAB8469S), p21$^{CIP1}$ (12D1, MAB2947) all from Cell Signaling Biotechnology, Beverly MA. Anti-rabbit and anti-mouse secondary antibodies were purchased from Cell Signaling Biotechnology Beverly, MA). Lipofectamine RNAimax were purchased from Thermofisher (Waltham, MA). BEAS2B (human bronchial epithelial cell line) (ATCC Teddington, UK) were cultured in keratinocyte media (Invitrogen, Paisley, UK) supplemented with human recombinant epithelial growth factor and bovine pituitary extract.

### Vesicles characterisation by nanoflow cytometry analysis

Large and small EVs were analyzed by NanoFCM (NanoFCM, Xiamen, China) for particle concentrations and size distribution. The instrument was calibrated for concentrations using polystyrene beads and for size distribution using Silica Nanosphere cocktail (NanoFCM Inc., S16M-Exo). Any particles that passed by the detector during a 1-min interval were recorded in each test. All samples were diluted in 0.22µm-filtered PBS to attain a particle count within the optimal range of 2000-12000/min. Using the calibration curve, EVs concentration and size were determined on the NanoFCM software (NanoFCM Profession V1.0).

Large and small EVs were treated for 20 min with 0.01% v/v Triton-X100 were used as an internal control.

### Western blotting

Protein extracts were prepared using RIPA buffer (Sigma: 150nM NaCl, 1.0% IGEPAL® CA-630, 0.5% (w/v) sodium deoxycholate, 0.1% (w/v) SDS, and 50mM Tris, pH 8.0) containing antiproteases (Roche, Welwyn Garden City, UK). Protein concentration of the isolated EVs samples were determined using the Pierce micro–BCA Protein Kit (ThermoFisher Scientific, Waltham, MA). Total protein lysates were mixed with NuPAGE LDS Sample (Invitrogen) and heated for 8 min, 110°C. Samples were loaded onto NuPAGE 4–12% (w/v) Bis-Tris Protein Gels (Invitrogen) and run with NuPAGE MOPS SDS Running (Invitrogen) according to the NuPAGE Novex electrophoresis program. Proteins were transferred to Nitrocellulose Blotting

Membranes (Invitrogen) and incubated cold overnight with primary antibody in blocking buffer. Next day, the membranes were washed and incubated with anti-mouse or anti-rabbit secondary antibody conjugated with HRP for 1 hour at room temperature. Western blot was analyzed by chemiluminescence (ECL Plus; GE Healthcare, Hatfield, UK) using an Odyssey CLx scanner and the ImageStudio Software (LI-COR Biosciences). Protein signals were normalized to β-actin signal.

**RNA extraction and real-time quantitative PCR**

mRNA and miRNAs were extracted using the miRNeasy kit (Qiagen) according to the manufacturer's instructions. RNAs and miRNA were reverse transcribed using the TaqMan total RNA including small RNA reverse Transcription kit (Applied Biosystems, Life Technologies). Both mRNA and miRNA levels were detected by either TaqMan Assays (SIRT1 Hs01009006, CDKN1A/p21 Hs00355782), or TaqMan MicroRNA Assay (hsa-miR-34a-5p 000426, hsa-miR-570-3p 002347) (Thermofisher). RNU48 (001006), a small noncoding RNA, was detected as the endogenous control for miRNA detection in cells and in EVs as its expression were not modified by any stimulation used in our study. Guanine nucleotide binding protein – polypeptide 2-like 1 (GNB2L1, encoding RACK1) was used as endogenous control for mRNA-derived cDNA. After the reactions, the CT values were determined using fixed-threshold settings. The relative fold difference was calculated using the $2^{-\Delta\Delta Ct}$ method.

**SA-β-galactosidase staining**

SAEC from healthy donors were plated into 24-well plates and left for 24h to adhere. Cells were treated with large EVs (~$5x10^{10}$ EVs) for 48h. Cells were then fixed, and senescence-associated β-galactosidase activity was determined according to the manufacturer's instructions (ab65351; Abcam).

**Online Data Supplement Figure 1**

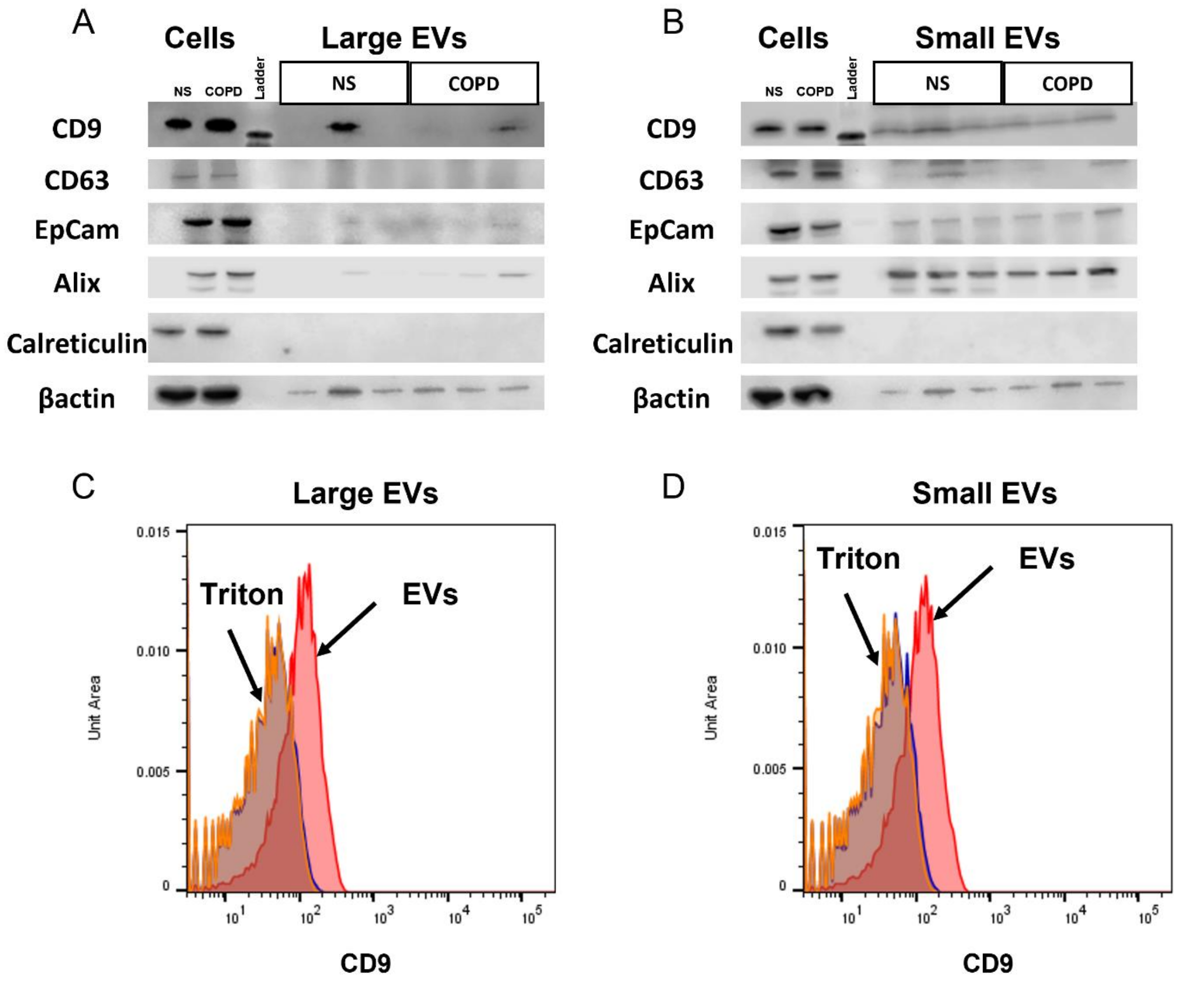


**Characterisation of EVs produced by SAEC**.

Large and small EVs were isolated from 7-days conditioned media from COPD or non-smoker SAEC. (A) Large EVs and (B) small EVs were resuspended in RIPA buffer and the expression of EVs marker was analysed by Western blot. Images are representative of 3 independent experiments. (C) Large EVs and (D) small EVs were treated with 0.01% of Triton X100 and analysed by flow cytometry using beads coated with CD9-antibody. Images are representative of 4 independent experiments.

**Online Data Supplement Figure 2**

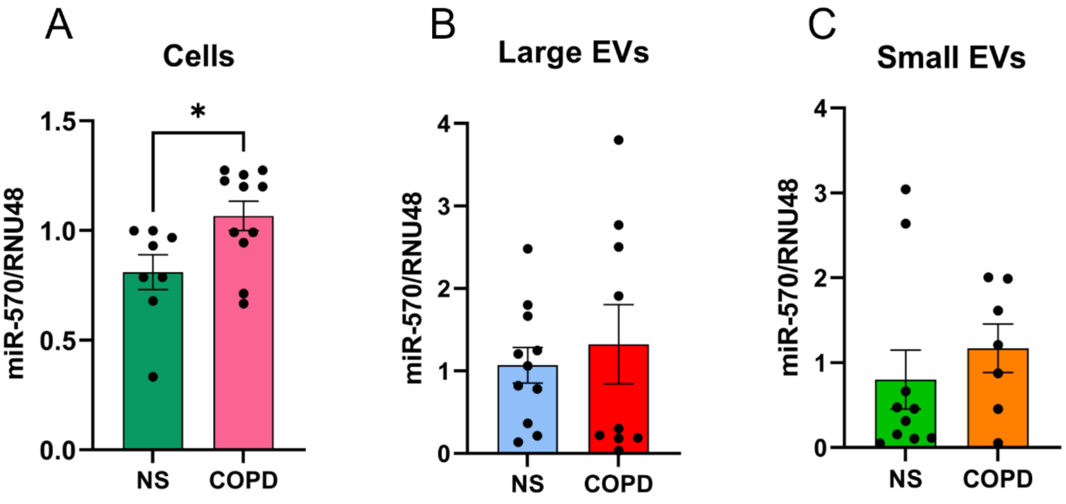


**EVs from COPD SAEC are loaded with miR-34a but not with miR-570**.

Large and small EVs produced by SAEC from healthy and COPD subjects were isolated from 7-days conditioned media. Total RNA was extracted from (a) cells (n=8), (b) large EVs (n=10) and (c) small EVs (n=10). MiR-570 expression was determined by RT-qPCR. Each point represents EVs isolated from different SAEC donors. Data are mean ± SEM, analysed by Mann-Whitney test.

**Online Data Supplement Figure 3**

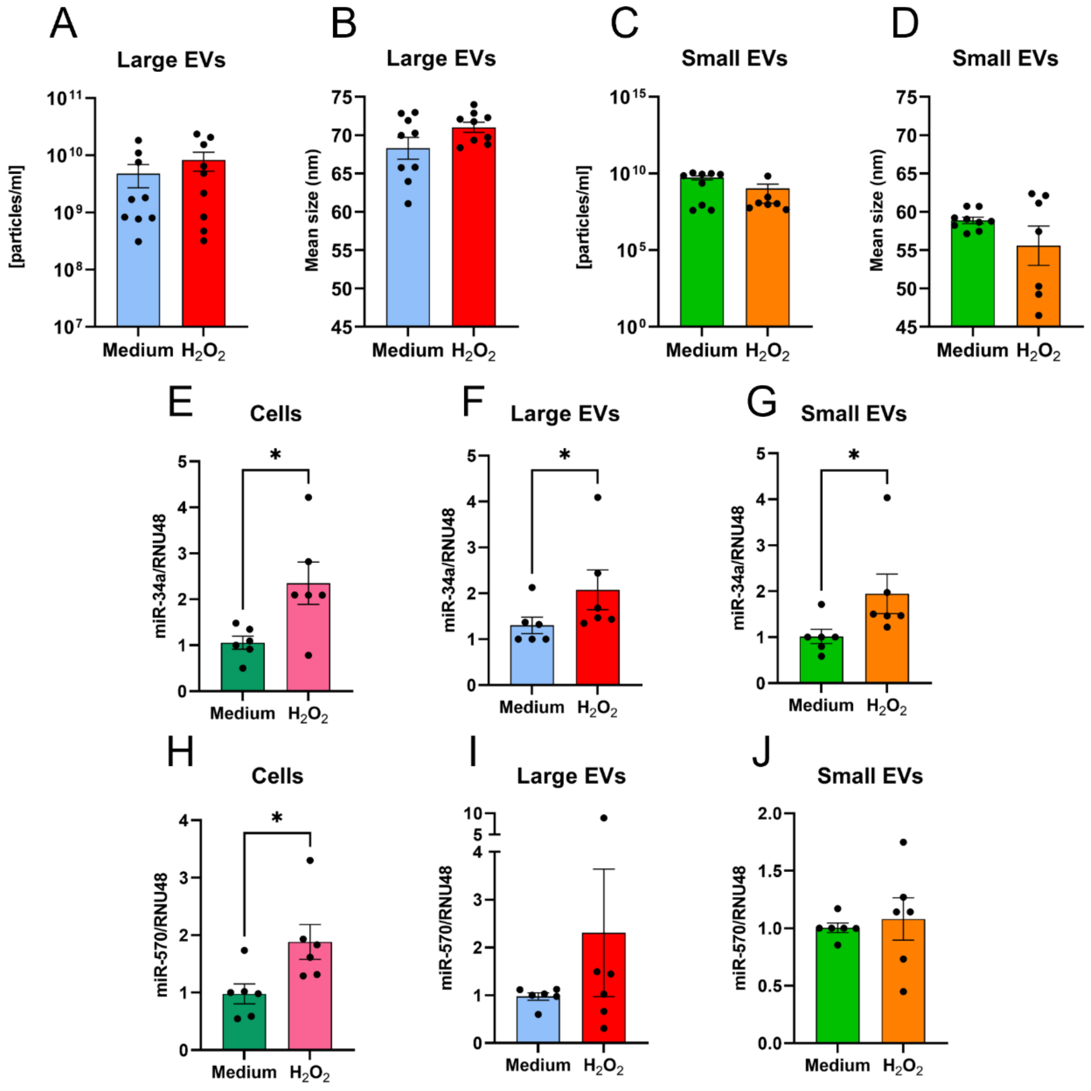


**In response to oxidative stress, BEAS2B cells produce extracellular vesicles enriched with miR-34a.**

BEAS2B cells were cultured for 48h with serum free medium or 100µM of hydrogen peroxide ($H_2O_2$) and EVs isolated from cell media. Concentration of EVs (**A and B**) and size of EVs (**C and D**) were determined by NanoFCM. Total RNA was extracted and changes in miR-34a expression (**E-G**) and miR-570 (**H-J**) were determined in cells (**E, H**, n=6), large EVs (**F, I**, n=6) and small EVs (**G, J**, n=5) by qRT-PCR. Data are mean ± SEM, analysed by Mann-Whitney test. *p<0.05

**Online Data Supplement Figure 4**

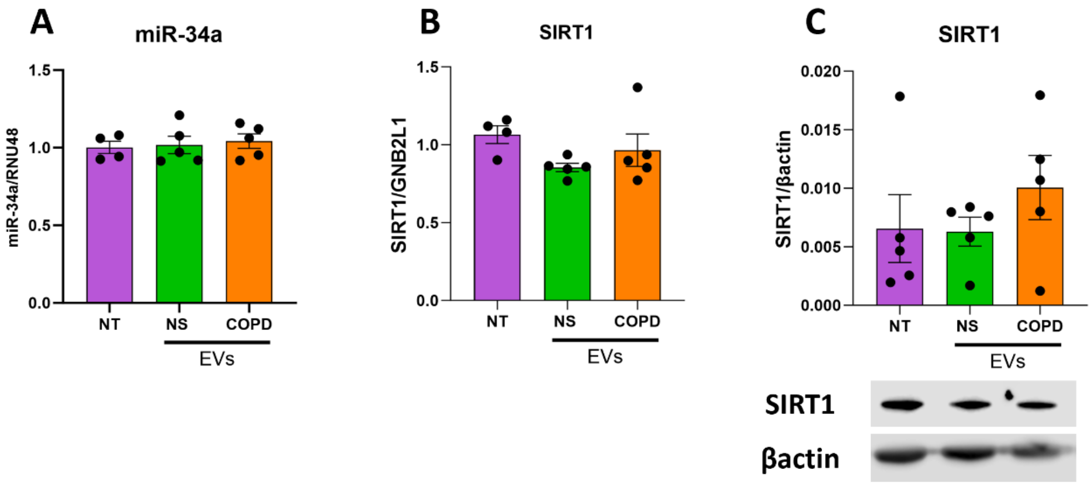


**Small EVs from COPD patients do not transfer miR-34a in recipient SAEC.**

Small EVs were isolated from 7-days conditioned media from COPD or non-smoker SAEC. 250 000 recipient healthy SAEC were stimulated with 150µl of small EVs and expression of (A) miR-34a, (B) SIRT1 were measured in recipient cells after 3h (for miR-34a) and 12h (SIRT1) by RT-qPCR. After 48h stimulation, (C) SIRT1 protein expression was analysed by western blotting. Each point represents EVs isolated from different SAEC donors (n=6). Data are expressed as mean ± SEM, analysed by Kruskal-Wallis test with post hoc Dunns or Mann-Whitney as appropriate where *p<0.05 ** p<0.01

**Online Data Supplement Figure 5**

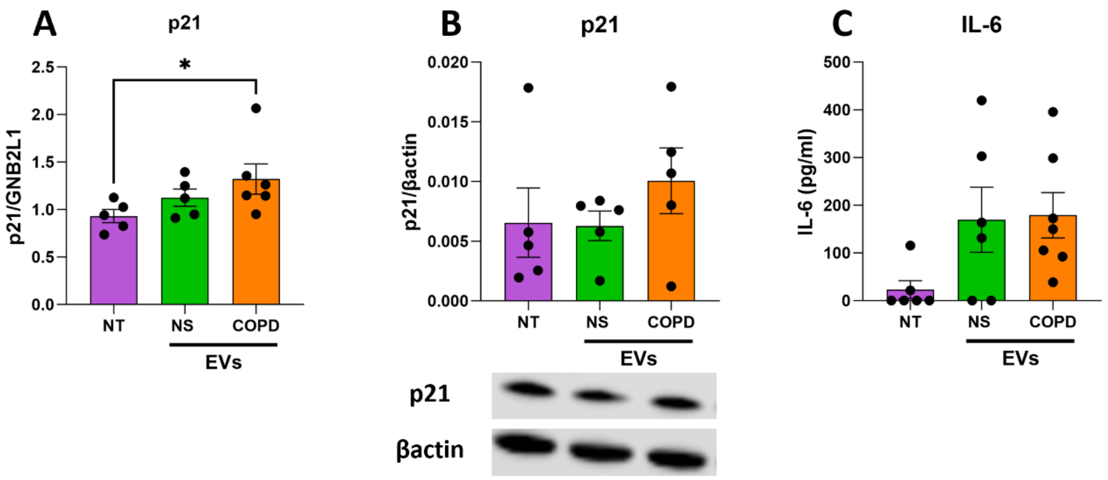


**Small EVs from COPD cells do not induce senescence markers in recipient SAEC.**

Small EVs were isolated from 7-days conditioned media from COPD or non-smoker SAEC. 250 000 recipient healthy SAEC were stimulated with 150µl of small EVs and expression of (A) p21 was measured in recipient cells and 12h. After 48h stimulation, (B) p21 protein was analysed by western blotting, and (C) IL-6 secretion was measured by ELISA. Each point represents EVs isolated from different SAEC donors (n=6). Data are expressed as mean ± SEM, analysed by Kruskal-Wallis test with post hoc Dunn's or Mann-Whitney as appropriate where *$p<0.05$ ** $p<0.01$

**Online Data Supplement Figure 6**

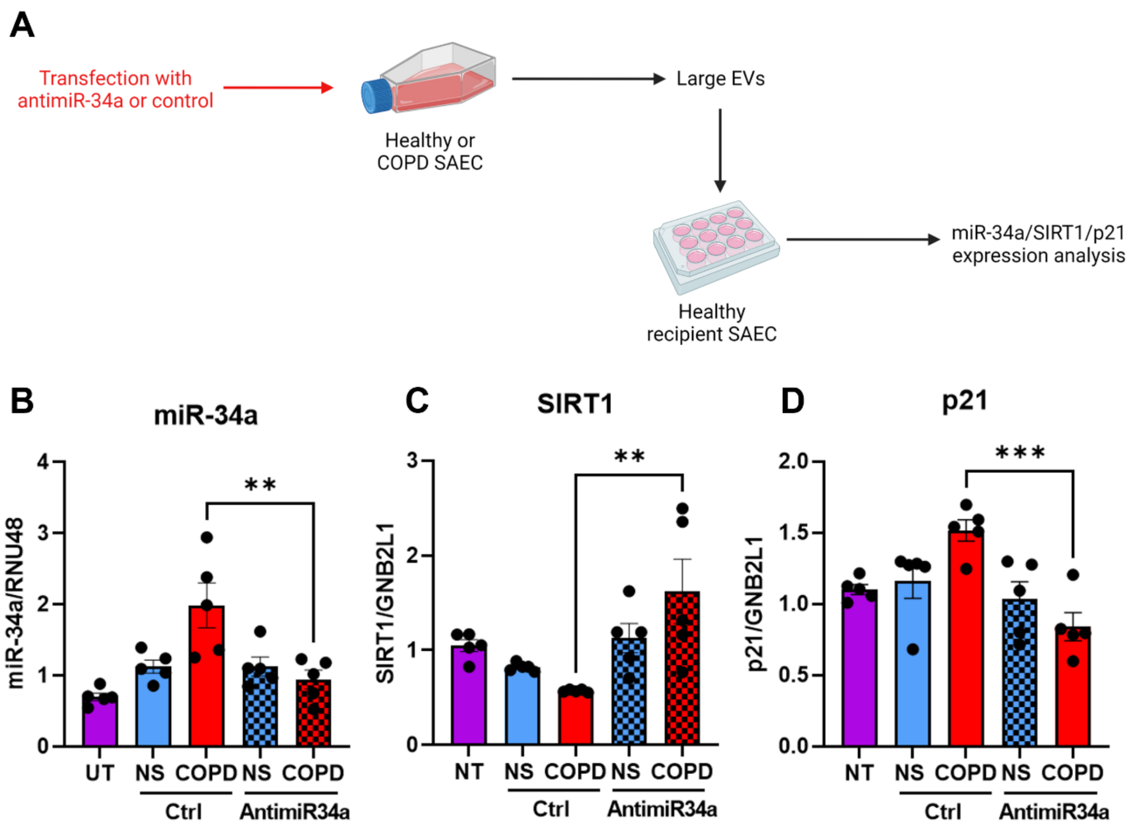


**The transfer of miR-34a loaded in EVs from COPD cells induce senescence in recipient cells**. (A) Donor healthy and COPD SAEC were transfected with an antagomir against miR-34a or with a control sequence and EVs were isolated from NS or COPD SAEC media. Healthy recipient SAEC were treated with EVs isolated from NS-ctrl, NS-antimiR-34a, COPD-ctrl or COPD-antimiR-34a. Changes in (B) miR-34a, (C) SIRT1 and (D) p21 mRNA were measured by qRT-PCR after 3 and 12 h stimulation. Each point represents treatment with EVs isolated from different SAEC donors. Data are expressed as mean ± SEM, analyzed by Wilcoxon test. **p<0.01, ***p<0.001
